# A new abundant nitrite-oxidizing phylum in oligotrophic marine sediments

**DOI:** 10.1101/2023.10.23.563599

**Authors:** Rui Zhao, Steffen L. Jørgensen, Andrew R. Babbin

## Abstract

Nitrite-oxidizing bacteria (NOB) are important nitrifiers whose activity regulates the availability of nitrite and links reduced ammonium and oxidized nitrate in ecosystems. In oxic marine sediments, ammonia-oxidizing archaea (AOA) and NOB together catalyze the oxidation of ammonium to nitrate, but the observed abundance ratios of AOA to canonical NOB are significantly higher than the theoretical ratio predicted from microbial physiology, indicating that many novel NOBs are yet to be discovered. Here we report a new bacterial phylum *Candidatus* Nitrosediminicolota, members of which are more abundant than canonical NOBs and are widespread across global oligotrophic sediments. *Ca.* Nitrosediminicolota members have the functional potential to oxidize nitrite, in addition to other accessory functions such as urea hydrolysis and thiosulfate reduction. While one recovered species (*Ca.* Nitrosediminicola aerophilis) is generally confined within the oxic zone, another (*Ca.* Nitrosediminicola anaerotolerans) can additionally thrive in anoxic sediments. Counting *Ca.* Nitrosediminicolota as a nitrite-oxidizer resolves the apparent abundance imbalance between AOA and NOB in oxic marine sediments, and thus its activity may exert a critical control on the nitrite budget.

## Introduction

Nitrite is an important intermediate compound in the biogeochemical nitrogen cycle, whose cycling dictates the availability of fixed nitrogen in marine ecosystems. Nitrite is controlled by multiple metabolic pathways: it can be produced by nitrate reduction and aerobic ammonia oxidation, and consumed by nitrite reduction and nitrite oxidation [reviewed in ^1^]. Among these pathways, by converting nitrite to nitrate, nitrite oxidation is a critical control point counteracting the further reduction of nitrite and subsequent fixed nitrogen loss to the atmosphere ^2^. Nitrite oxidation is mediated by a phylogenetically diverse functional guild known as the nitrite-oxidizing bacteria (NOB), which has been studied in a range of ecosystems, such as engineered environments ^3, 4^, coastal sediments ^5, 6^, haloalkaline lake sediments ^7^, hot springs ^8, 9, 10^, seawater ^11, 12^, and oxygen deficient zones ^13^. However, the diversity and metabolic capacities of NOB in deep-sea sediments have not been well studied.

Nitrification is catalyzed by two different chemolithoautotrophic guilds, ammonia oxidizers and nitrite oxidizers, and is an important nitrogen cycling process in global marine sediments. Nitrifiers seem to be dominant among the microbial communities in oxic sediments ^14, 15^, which account for a considerable proportion of the global seafloor ^16^. Ammonia-oxidizing archaea (AOA) are well known to dominate over ammonia-oxidizing bacteria in the marine environment, including sediments^14, 17, 18^, and have been well studied ^15, 19, 20^. By comparison, our knowledge about microorganisms involved in nitrite oxidation in marine sediments is extremely limited. Previously, gene-based surveys have indicated the presence of *Nitrospinaceae* and *Nitrospiraceae* in marine sediments ^14, 18, 21^, with a handful of cultured representatives from coastal sediments ^5, 6^. However, it remains unclear whether members of *Nitrospinaceae* and *Nitrospiraceae* (i.e., the canonical marine nitrite oxidizers) are in fact the major NOBs in marine sediments.

Nitrite rarely accumulates in marine sediments ^22, 23^, especially in the oxic zone where nitrite is mainly produced by ammonia oxidation. Due to the presence of oxygen, newly produced nitrite is rapidly oxidized to nitrate by NOB. The absence of appreciable nitrite (and ammonium) in this zone also indicates that NOBs are as efficient in the oxidation of nitrite as AOA in the oxidation of ammonium released from organic matter degradation. When grown in ocean-relevant conditions, marine AOA exhibit approximately 2.3-fold higher biomass yields (per nitrogen oxidized) than NOB but maintain only 1/3 the cell quota of NOB ^24^. In oxic sediments without nitrite accumulation, because of the balanced bulk reaction rates of ammonia and nitrite oxidation, the cell abundance of AOA should be theoretically 6.9 times higher than that of NOB. However, the observed abundances of canonical marine NOB are always orders of magnitude lower than those of AOA in water columns and marine sediments [e.g., ^14, 18, 25, 26, 27^]. The abundance discrepancy suggests that the majority of NOB are yet to be identified, likely because of limited studies and high diversity. Alternatively, if AOA severely outnumber NOB in oxic marine sediments, the nitrogen cycle might not be closed and there must be a cryptic nitrite loss process, presumably via denitrification that consumes bio-available nitrogen rather than recycling it ^2^.

In this study, we first highlight an abundance mismatch between AOA and canonical NOB in oxic sediments, based on a compilation of quantitative data in 11 marine sediment cores. To resolve this discrepancy, we rely on metagenome sequencing data from the Arctic Mid-Ocean Ridge to discover overlooked NOBs. We focus on three metagenome-assembled genomes (MAGs) that contain the metabolic potential of nitrite oxidation and form a bacterial phylum different from previously known NOB phyla. We then search for the presence of these novel NOB across global marine sediments. We conclude by calculating the abundance ratio of AOA to NOB with this new, more abundant phylum to resolve the previously identified discrepancy.

## Results and Discussion

### Abundance mismatch between AOA and canonical NOB in oxic marine sediments

We tested whether the expected AOA to NOB abundance ratio of 6.9:1 is observed in oxic marine sediments by comparing the abundances of AOA and NOB in the oxic zones of 11 sediment cores. The sediment cores considered include eight from the Arctic Mid-Ocean Ridge (AMOR) [four cores reported in ^28^, plus GS13-CC2 ^29^, GS14-GC04 ^22^, GS14-GC02, and GS15-GC01 ^30^], one piston core (NP-U1383E ^14, 31^) retrieved from the North Pond of the Mid-Atlantic Ridge, and two piston cores from the North Atlantic Gyre ^15^. Oxygen in all but one (GS14-GC04) core penetrates at least 50 cm below the seafloor ^14, 15, 28^. The thick oxic zones permitted high-resolution profiles of molecular biology and geochemistry across the oxygenated sediments.

By initially assuming that only members of the canonical marine NOB families *Nitrospinaceae* and *Nitrospiraceae* can perform nitrite oxidation, we observe that the abundance of these NOB in individual cores is at least one order of magnitude lower than that of AOA at most investigated depths (Fig. S1). When combining the 11 sediment cores (a total of 82 samples) (Fig. 1A), we observe that the AOA:NOB abundance ratio is higher than the theoretical 6.9:1 in 71 of 80 samples (median = 43.3, with the 99% confidence interval [16.6, 83.9]; Fig. 1C). This apparent discrepancy between AOA and NOB abundances most likely indicates that there are many yet unidentified NOB present in marine sediments.

**Fig. 1.**
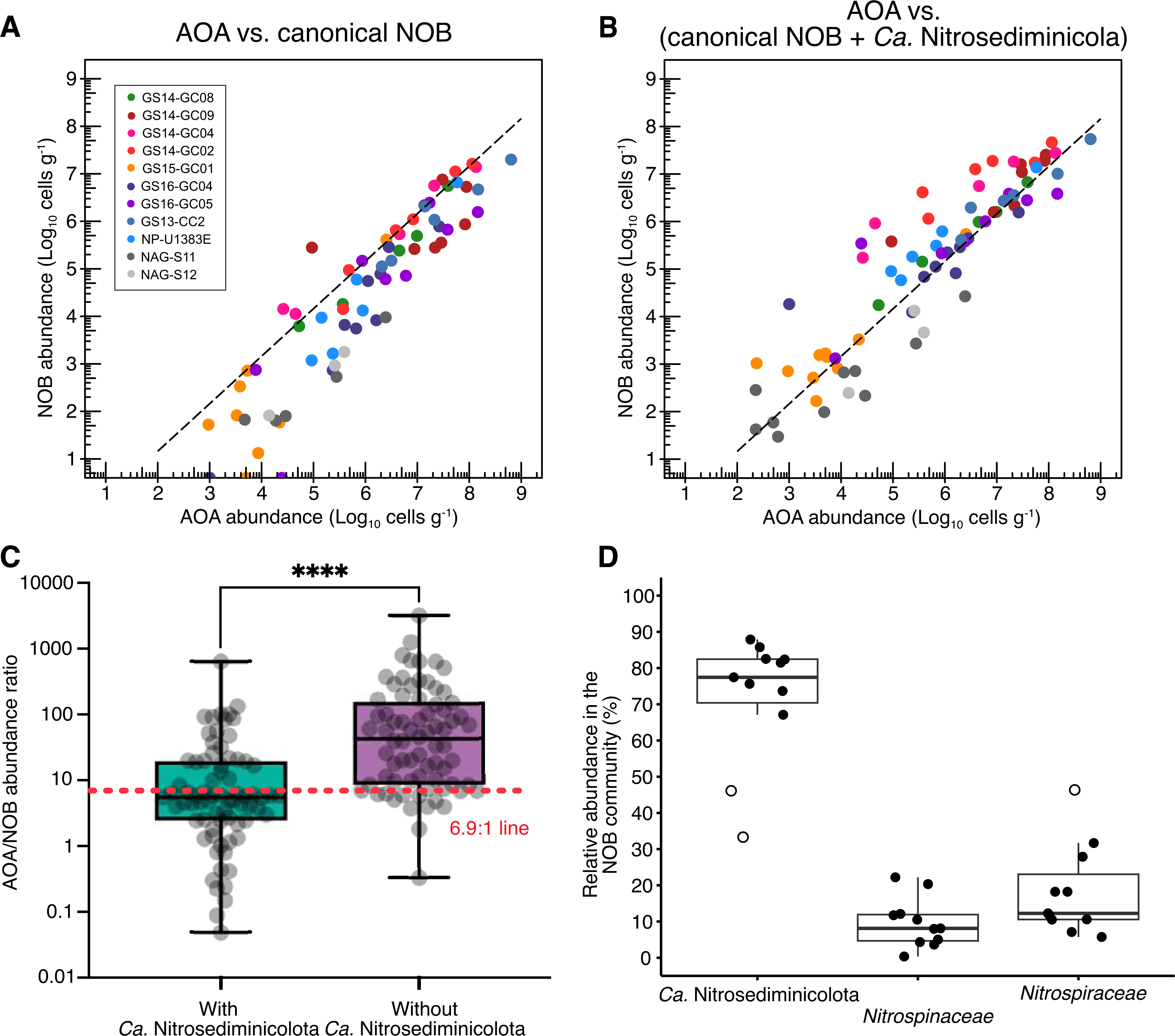
Comparison of the abundances of ammonia-oxidizing archaea (AOA) and nitrite-oxidizing bacteria (NOB) in oxic marine sediments. **(A)** Abundances of AOA and canonical NOB (affiliated with *Nitrospiraceae* and *Nitrospinaceae*) in a total of 82 samples of 11 sediment cores with extensive oxic zones. **(B)** Same as **A**, but AOA vs NOB including *Candidatus* Nitrosediminicolota. **(C)** Abundance ratios of AOA to NOB with and without *Ca.* Nitrosediminicolota in the total 82 samples. The colored boxes show the 99% confidence intervals. To facilitate the comparison, a dashed line delineating an AOA:NOB ratio of 6.9:1 is added in panels **A**, **B**, and **C**. (**D**) Depth-integrated relative abundances of the three NOB lineages in the oxic zones of the 11 investigated sediment cores. Boxes indicate 95% confidence intervals. Outliers are marked with open circles.

### A new bacterial phylum defined by MAGs from marine sediments

To elucidate which microbes are likely overlooked NOBs in marine sediments, we focused on the metagenome sequencing data generated for two investigated sediment locations: four sediment horizons of AMOR core GS14-GC08, four horizons of NP-U1383E of North Pond ^28^. Through genome binning and refinement, we noticed three MAGs (Bin_086, Bin_096, and Bin_108) that contain genes encoding the nitrite-oxidizing enzyme nitrite oxidoreductase (Nxr) but are not affiliated with any well-defined bacterial phylum. All three MAGs are of high completeness (> 92%; Table 1) and low fragmentation (<73 scaffolds; Table 1) and therefore should be regarded as high-quality genomes. The genome sizes are in the range of 1.8–2.4 Mbp. Automatic classification based on the 120 bacterial single-copy genes suggests that they are affiliated with an understudied bacteria phylum (with the placeholder JADFOP01 in the GTDB RS214 Release), which previously included three MAGs (B6D1T2, B58T1B8, and B13D1T1) recovered from hadal sediments beneath the Mariana Trench ^32^.

The novel phylogenetic affiliation of the now total six MAGs included in the JADFOP01 phylum is confirmed by phylogenetic analyses. Within the phylogenetic trees based on the concatenated 120 bacterial single-copy genes (Fig. 2B) and 14 conservative single-copy ribosomal proteins (Fig. S2), the six MAGs form a branch separated from several established bacterial phyla such as Nitrospinota (containing NOB), Tectomicrobia, Nitrospinota_B, Schekmanbacteria, and UBA8248 (Fig. 2A). The average nucleotide identities (ANI) between members of JADFOP01 and those in the established phyla are in the range of 47–53%, much lower than the threshold of 83% distinguishing bacterial phyla ^33^ and supporting the view that these MAGs represent a new bacterial phylum. The novel phylogenetic affiliations of these MAGs are confirmed by phylogenetic analysis based on the 16S rRNA gene, which shows a congruent topology with that based on both 120 single-copy genes and ribosomal proteins (Fig. 2B). We tentatively name this phylum *Candidatus* Nitrosediminicolota, for their prevalence in globally distributed marine sediments (see Etymology description).

**Fig. 2.**
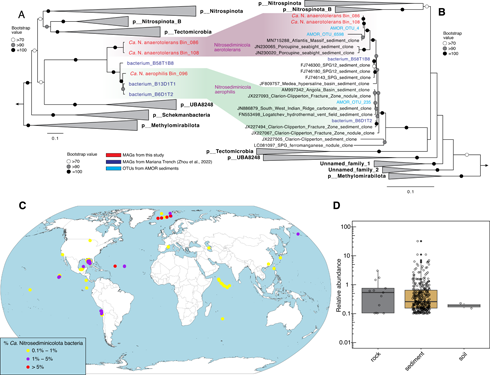
Phylogeny and distribution of *Candidatus* Nitrosediminicolota. **(A)** Maximum-likelihood phylogenetic tree of *Ca.* Nitrosediminicolota and related phyla based on the 16S rRNA gene. **(B)** As **A**, but the maximum-likelihood phylogenetic tree of *Ca.* Nitrosediminicolota and related phyla based on the concatenated 120 single-copy genes of bacteria. Both trees are rooted in five Methylamirales genomes. The two MAGs recovered from AMOR sediments are highlighted in red. The nomenclature of the bacterial phyla follows GTDB, except that *Ca.* Nitrosediminicolota was proposed in this study. Bootstrap values of >70 (*n* = 1000) are shown with symbols listed in the legend. The scale bars show estimated sequence substitutions per residue. **(C)** Global distribution of *Ca.* Nitrosediminicola bacteria. Except for two soil sites and two basaltic rock sites, *Ca.* Nitrosediminicola bacteria are present in multiple depths of each of the sediment cores represented by individual circles. In each core, the maximum relative abundance is shown using different colors as listed in the legend. (**D**) Relative abundances of *Ca.* Nitrosediminicolota bacteria in three major habitats where they were detected at >0.1% relative abundance.

The calculated average amino acid identities (AAI) between the six MAGs are >80% (Fig. S3), placing them in the range suggested for genomes belonging to the same genus [65–95%, ^33^] and therefore indicating that they should belong to a single genus. We tentatively name this genus *Candidatus* Nitrosediminicola. Within this genus, three MAGs (Bin_096, B6D1T2, and B13D1T1) show AAI higher than 95% and should fall into the same species, for which we suggest a provisional name *Candidatus* Nitrosediminicola aerophilis. Bin_086 and Bin_108 also shared AAI higher than 95% and belong to the same species, which we provisionally name *Candidatus* Nitrosediminicola anaerotolerans. The remaining MAG B58T1B8 shows AAI of <79% with all other *Ca.* Nitrosediminicola MAGs and therefore should represent a third species that we do not name as it is not found in our samples. Therefore, the six MAGs of *Ca.* Nitrosediminicola reported here should be resolved to three species within a single genus.

### *Ca.* Nitrosediminicolota are prevalent in oligotrophic marine sediments

To explore semi-quantitatively the global occurrence of *Ca.* Nitrosediminicolota, we searched public amplicon sequencing datasets in the IMNGS database ^34^ for the 16S rRNA gene sequences of our high-quality MAGs (See Materials and Methods for details). *Ca.* Nitrosediminicolota is present with >0.1% relative abundances in 300 globally-distributed samples, which are mapped in Fig. 2C. Except for two soil and 13 basaltic rock samples (from the Dorado outcrop ^35^ and North Pond ^36^), the vast majority of the *Ca.* Nitrosediminicolota-containing samples are marine sediments (Fig. 2D). All of the marine sites are oligotrophic sediments beneath the oligotrophic gyres of the Pacific ^37^, Atlantic ^15^, and Indian Oceans ^38^, mid-ocean ridges ^22, 28, 30^, hadal trenches ^39, 40^, and the Gulf of Mexico ^41^ (Fig. 2C). The distribution of the *Ca.* Nitrosediminicolota phylum suggests it harbors microbes specialized for oligotrophic marine sediments.

### *Ca.* Nitrosediminicolota contain all key machinery of nitrite oxidizers

*Ca.* Nitrosediminicolota members contain nitrite oxidoreductase (NXR), the key enzyme for nitrite oxidation in microorganisms. NXR is present in four *Ca.* Nitrosediminicola MAGs (Bin_086, Bin_108, B6D1T2, and B58T1B8) that span all three species in this genus (Fig. 3A). Considering the high similarities (>95% AAI) among the three MAGs represented by B6D1T2, it is likely that the absence of NXR in the other MAGs (Bin_096 and B13D1T1) of this species is due to their lower genome completeness (Table 1). The continuous NXR operon within *Ca.* Nitrosediminicolota genomes, consisting of NxrABC and a chaperone subunit which annotates as NxrD, may enable members of *Ca.* Nitrosediminicola to generate energy from the oxidation of nitrite.

**Fig. 3.**
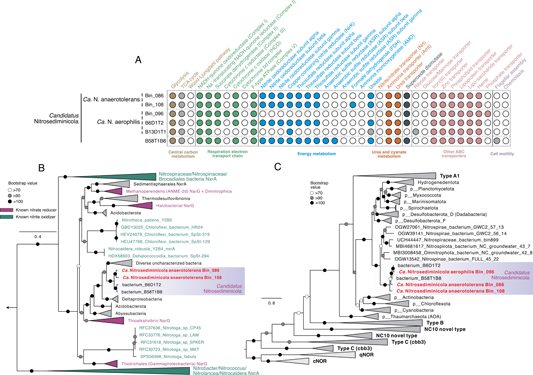
Metabolic potential of *Ca.* Nitrosediminicolota bacteria. **(A)** Heatmap showing the important metabolic pathways encoded by the six *Ca.* Nitrosediminicolota genomes. The filled circles indicate the presence of the full pathways, the open ones denote the absence, while the grey ones represent that the pathways are incomplete. **(B)** Phylogeny of NxrA/NarG of the novel NOB. The tree is rooted to NarG sequences of NC10 bacteria. Genomes recovered in this study are shown in red. Bacteria known for having the capacity of nitrite oxidation (i.e., nitrite-oxidizing bacteria of the genera of *Nitrospira*, *Nitrospina*, *Nitrotoga*, *Nitrobacter*, and *Nitrococcus*, and anammox bacteria of the Brocadiales order) are highlighted in green. Bacteria with an observed nitrate-reducing phenotype are shown in purple. **(C)** Maximum-likelihood phylogenetic tree of heme copper reductase (or cytochrome *c* oxidase). The sequences of *Candidatus* Nitrosediminicolota are highlighted in a colored box, and the MAGs recovered from AMOR sediments are shown in red. For both trees, bootstrap values of >70 (*n* = 1000) are shown with symbols listed in the legend. The scale bar shows estimated sequence substitutions per residue.

Aerobic NOBs need oxygen as their terminal electron acceptor. Five of the six *Ca.* Nitrosediminicola genomes contain a cytochrome *c* oxidase (CoxABCDE) (i.e., heme-copper oxygen (HCO) reductase) (Fig. 3A), a critical enzyme involved in oxygen respiration. Phylogenetic analysis of cytochrome *c* oxidase indicates that the sequences of *Ca.* Nitrosediminicola form a clade separated from other bacterial phyla and fall within the broad branch of the A1 Clade of heme-copper oxygen reductase (Fig. 3C). *Ca.* Nitrosediminicola members lack the cytochrome *bd*-type oxidases that are common in *Nitrospinaceae* ^5, 42^ and *Nitrospiraceae* ^43^ or *cbb3*-type cytochrome *c* oxidase. The cytochrome *c* oxidase can receive electrons from NXR for aerobic respiration, and the protons released by this process can help to maintain the proton gradient that drives the ATP synthesis in Complex V. The presence of oxygen-respiring cytochrome *c* oxidase likely also enables them to complete the electron-transport chain and support the high abundances of *Ca.* Nitrosediminicola in oxic sediments.

Characterized NOBs have been suggested to acquire the NXR module in different evolutionary pathways and the horizontal transfer of NXR is likely a major driver for the spread of the capability to gain energy from nitrite oxidation during bacterial evolution ^8, 10, 42, 44, 45^. In particular, the canonical marine aerobic NOBs affiliated to the genera *Nitrospira* and *Nitrospina* are suggested to obtain their NXR from anammox bacteria in the Brocadiales order within the Planctomycetota phylum ^29^. Considering the distinct phylogenetic affiliations between the newly found *Ca.* Nitrosediminicola and the canonical NOB, we checked whether they acquired the nitrite oxidation capacity through the same evolutionary path. Phylogenetic analysis of the NXR alpha subunit (NxrA) suggested that the four NXR-bearing *Ca.* Nitrosediminicola members fall into the so-called “new NXR/NAR clade” defined by ^45^, which includes some recently characterized NOB such as *Nitrotoga* ^45, 46, 47^ and Chloroflexota NOBs (*Ca.* Nitrocaldera robusta and *Ca.* Nitrotheca patiens ^10^), rather than the *Nitrospiraceae*/*Nitrospinaceae*/Anammox clade. (Fig. 4). *Ca.* Nitrosediminicolota members thus may have acquired NXR from a donor different from those taxa within the *Nitrospiraceae*/*Nitrospinaceae*/Anammox clade.

**Fig. 4.**
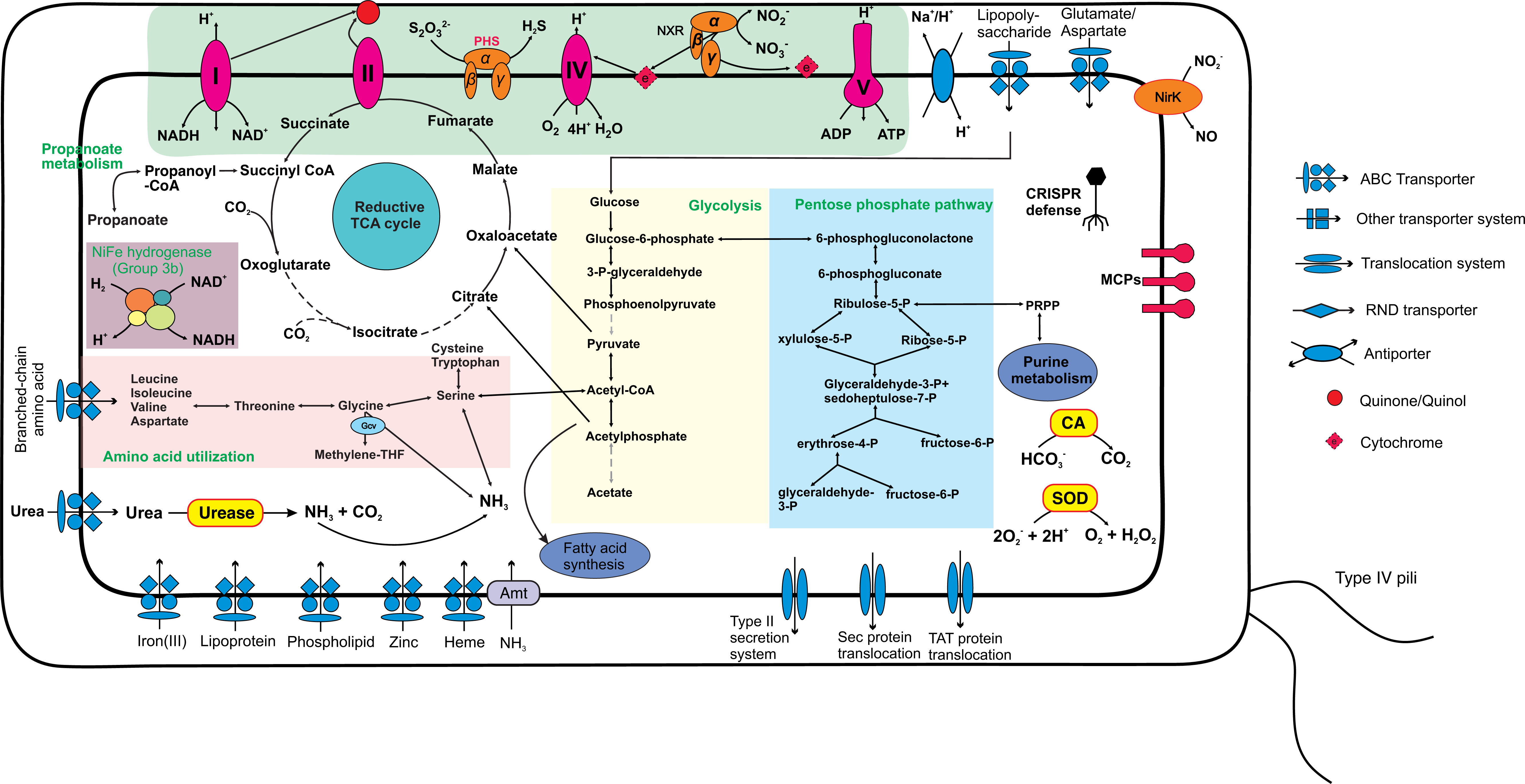
Potential key metabolic interactions in *Ca.* Nitrosediminicolota bacteria. The common metabolic pathways in the *Ca.* Nitrosediminicolota bacteria include aerobic respiration, nitrite oxidation (NXR), oxygen respiration (Complex IV), urea assimilation and hydrolysis, reductive TCA cycle, nitrite reduction (NirK), glycolysis, superoxide dismutase (SOD), pentose phosphate pathway, and ABC transport for iron, zinc, heme, lipoprotein, and phospholipid.

*Ca.* Nitrosediminicola members can also reduce nitrite via nitrifier denitrification like other NOB ^48^. All but one (Bin_096) of the *Ca.* Nitrosediminicola genomes encode a copper-containing nitrite reductase (NirK), which can reduce nitrite to nitric oxide (Fig. 3A) and is common among NOBs ^6, 44, 49^. On the maximum-likelihood phylogenetic tree of bacterial NirK *Ca.* Nitrosediminicola genomes form an independent cluster (Fig. S4). The close relatives of *Ca.* Nitrosediminicola NirK are all from ultra-small-celled archaea affiliated with *Ca.* Woesearchaeota, rather than Nitrospinota, Nitrospinota_B, Tectomicrobia, UBA8284, or Schekmanbacteria, indicating that NirK in *Ca.* Nitrosediminicola may have a different origin than the majority of the *Ca.* Nitrosediminicola genes.

For the full respiratory electron-transport chain, *Ca.* Nitrosediminicola genomes have Complex I, Complex II, Alternative Complex III, Complex IV (described above), and Complex V (F-type ATPase) (Fig. 3A), like other previously characterized aerobic NOB. This complete oxygen respiratory electron-transport chain likely enables them to oxidize nitrite under oxic conditions. Regarding the central carbon metabolism, similar to the two newly-cultured NOBs affiliated with *Nitrospinaceae* from coastal sediments ^5^, *Ca.* Nitrosediminicola species encode an (incomplete) tricarboxylic acid (TCA) cycle (Fig. 3 and 4), which may be used to fix CO_2_ as proposed previously for *Nitrospira* and *Nitrospina* ^42, 44, 50^. The electrons for carbon fixation may be derived from nitrite oxidation ^42^. The *Ca.* Nitrosediminicola genomes also encode the gluconeogenesis and the pentose phosphate pathways (Fig. 4), which may be employed for the synthesis of precursor metabolites in these NOBs, similar to that previously proposed for *Nitrospira moscoviensis* ^50^. Another feature of *Ca.* Nitrosediminicola is that five of its six member genomes contain a urease operon (Fig. 3A) (See Supplementary Text), which may enable them to access this pool for substrates and energy and also engage in reciprocal feeding with co-occurring ammonia-oxidizing archaea ^14, 19^ to increase their metabolic fitness in marine sediments.

### *Ca.* Nitrosediminicolota resolves the nitrifier abundance discrepancy

Given the likely nitrite oxidation capacity of the newly defined *Ca.* Nitrosediminicolota, we re-calculated the abundances of combined NOB (defined as the sum of *Nitrospiraceae*, *Nitrospinaceae*, and *Ca.* Nitrosediminicolota) in the 11 sediment cores by including these as part of the NOB community. The abundances of NOB and AOA in the oxic zones falls closer to the 6.9:1 line (Fig. 1B). The median of AOA:NOB ratio of the total 82 oxic samples decreases dramatically from 43.3 to 5.6, with a 99% confidence interval of 3.9–8.4 (Fig. 1C). These results align with the theoretical prediction based on the observed features of marine AOA and NOB. Therefore, counting these novel bacteria as NOBs helps resolve the apparent abundance mismatch between AOA and NOB in marine sediments.

To check whether *Ca.* Nitrosediminicolota is the dominant nitrite oxidizer in both oxic and anoxic marine sediments, we compared the abundances of *Ca.* Nitrosediminicolota to those of canonical NOB affiliated with the families *Nitrospinaceae* and *Nitrospiraceae* in AMOR sediment cores. Our results suggest that *Nitrospinaceae* and *Nitrospiraceae* are generally confined within the oxic zone with <4% relative abundances among the total prokaryotic communities (Fig. 5B and Fig. S6B), probably because these NOBs require oxygen as the electron acceptor for nitrite oxidation and cannot tolerate strict anaerobic conditions. In contrast, *Ca.* Nitrosediminicolota is present in most of the investigated depths and is particularly abundant in the deep anoxic layers. Restricting analysis to the oxic zone where the organisms co-occur, *Ca.* Nitrosediminicolota dominates over *Nitrospinaceae* and *Nitrospiraceae* in all but a few depths (Fig. 5D and Fig. S6D). The depth-averaged relative abundance of *Ca.* Nitrosediminicolota in the putative NOB communities in oxic zones of the 11 cores is calculated to be 50–80%, while *Nitrospiraceae* and *Nitrospinaceae* each only accounts for 8–25% (Fig. 1D). Thus, *Ca.* Nitrosediminicolota is roughly 2–4 times more abundant than the canonical NOBs in oxic marine sediments. The dominance of *Ca.* Nitrosediminicolota is also evident in the calculated absolute abundance profiles in the four AMOR cores (Fig. 5C and Fig. S6C), who exhibit 2–4 orders of magnitude higher absolute abundances than *Nitrospinaceae* and *Nitrospiraceae* in the basal part of the oxic zones and also anoxic sediment layers. Being far more abundant than canonical NOBs, *Ca.* Nitrosediminicolota can contribute significantly to nitrite oxidation in global oligotrophic marine sediments.

**Fig. 5.**
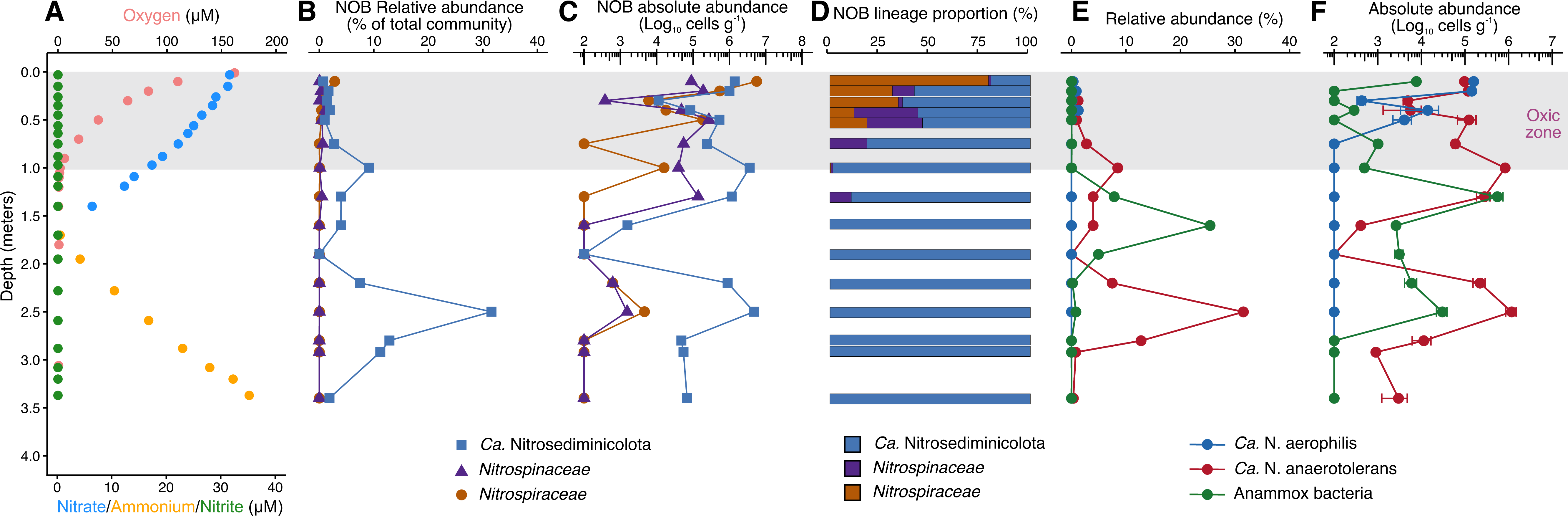
Geochemical context, relative abundances, and community compositions of NOB lineages in AMOR core GS14-GC08. **(A)** Geochemical context delineated by the measured profiles of oxygen, nitrate, nitrite, and ammonium, previously reported in ^28^. The oxic zone is marked with a grey box. **(B)** The relative abundances of *Ca.* Nitrosediminicolota and the canonical marine NOB families *Nitrospiraceae* and *Nitrospinaceae*, as assessed by amplicon sequencing. **(C)** The absolute abundances of the three NOB lineages calculated as the product of the relative abundances of the three lineages and the total cell numbers. (**D**) The community composition of NOB community in each of the investigated depth. (**E, F**) The relative (**E**) and absolute (**F**) abundances of two *Ca.* Nitrosediminicola species (*Ca.* N. aerophilis and *Ca.* N. anaerotolerans) and anammox bacteria throughout the core. The same data for other three AMOR cores are shown in Fig. S6.

### Redox niches distinguish *Ca.* Nitrosediminicola species

To reveal what *Ca.* Nitrosediminicolota species are present and can thrive in the anoxic sediment layers of the AMOR cores, we interrogated the distribution of individual species represented by the *Ca.* Nitrosediminicola MAGs in the four AMOR cores previously reported ^28^. Based on the comparison of 16S rRNA gene sequences between *Ca.* Nitrosediminicola MAGs and the amplicon sequencing OTUs (See Materials and Methods), *Ca*.

Nitrosediminicola Bin_086 and *Ca.* Nitrosediminicola Bin_096 reported here correspond to OTU_4 and OTU_235 reported in ^28^, respectively. While these two OTUs were previously classified as members of the Schekmanbacteria phylum (SILVA 138.1) or the Nitrospinota phylum (MD2896-B214 class, SILVA 128), they are in fact members of the newly established Nitrosediminicolota phylum. The matches between the MAGs reported here and the previously reported OTUs set the basis for tracking the vertical distribution of the two *Ca.* Nitrosediminicola species in the four AMOR cores.

We observed distinct redox niche preferences of the two *Ca.* Nitrosediminicola species derived from the AMOR cores. While *Ca.* Nitrosediminicolota Bin_096 (OTU_235) is exclusively detected in the oxic zones of the four AMOR cores, *Ca.* Nitrosediminicolota Bin_086 (OTU_4) was detected in all investigated sediment layers and is particularly abundant in anoxic layers (Fig. 5E and Fig. S6E). Therefore, *Ca.* Nitrosediminicolota Bin_096 appears to be an oxic niche specialist, while *Ca.* Nitrosediminicolota Bin_086 may be an anoxia-tolerant generalist. The high relative abundance of *Ca.* Nitrosediminicolota in the deep anoxic layers of the AMOR cores (Fig. 5B and Fig. S6B) is due to the prevalence of *Ca.* Nitrosediminicola Bin_086 in the anoxic sediments. The maximum relative abundance of *Ca.* Nitrosediminicola Bin_086 (31% of the total community) was detected at 250 cm below seafloor of GS14-GC08 (Fig. 8B). Such a redox niche preference difference between different lineages of the same functional guild is not novel, but has been previously observed for anammox bacterial families in AMOR sediments ^22^. To reflect their preferred redox niches, we propose to name Bin_096 *Ca.* Nitrosediminicola aerophilis (prefer aerobic conditions), and Bin_086 *Ca.* Nitrosediminicola anaerotolerans (tolerant to anaerobic conditions).

Because *Ca.* N. anaerotolerans is abundant in anoxic layers and can consume nitrite, they might also compete for nitrite with anammox in the same depths. In GS14-GC08, the relative and absolute abundances of *Ca.* N. anaerotolerans exhibited two peaks in the anoxic sediments (Fig. 5E and 5F), which are located above and below the nitrate-depletion zone. Interestingly, the peak (of both the relative and absolute abundances) of anammox bacteria was observed between the two *Ca.* N. anaerotolerans abundance maxima, which appears to indicate potential competition between these two nitrite-consuming groups. Although the two abundance peaks of *Ca.* N. anaerotolerans are not resolved in the other AMOR cores (GS14-GC09, GS16-GC04, and GS16-GC05) due to their short core length or low depth resolution, they generally show that the abundance of *Ca.* N. anaerotolerans decreases along with the increase of anammox bacteria abundance in anoxic sediments (Fig. S6). This observation suggests that anammox bacteria retain better fitness over *Ca.* N. anaerotolerans in anoxic sediments where nitrate and nitrite are limiting.

To identify what mechanisms may drive the distinct redox niche preferences between the two prevailing Nitrosediminicolota species, we performed a comparative genomic analysis based on four MAGs (*Ca.* N. anaerotolerans represented by Bin_086 and Bin_108, and *Ca.* N. aerophilis represented by Bin_096 and B6D1T2). A total of 5,928 genes of the four MAGs form 1,356 gene clusters. There are 47 gene clusters uniquely present in *Ca.* N. anaerotolerans and 37 in *Ca.* N. aerophilis. Thiosulfate reductase is among the enzymes encoded by the genes uniquely present in *Ca.* N. anaerotolerans. Thiosulfate is a common sulfur cycle intermediate at low concentrations in marine sediments and is mainly produced by the oxidation of hydrogen sulfide derived from sulfate reduction ^51, 52^. The presence of thiosulfate reductase in *Ca.* N. anaerotolerans may equip it with the capacity to use thiosulfate as an electron acceptor in anoxic sediments. In addition, *Ca.* N. anaerotolerans uniquely encode genes in ABC-type amino acid transport, cysteine protease, and lactate dehydrogenase, which together may enable *Ca.* N. anaerotolerans to assimilate, degrade, and ferment amino acids for energy conservation. Finally, unlike *Ca.* N. aerophilis, *Ca.* N. anaerotolerans also contains genes involved in cobalamin (Vitamin B12) synthase, which may be helpful for them to grow high density (>10^6^ cells g^-1^; Fig. 5C) in the anoxic sediments.

## Conclusion

Our compilation of AOA and canonical NOB abundances in global oxic marine sediments argues that there were overlooked yet abundant NOB. Through genome reconstruction and phylogenetic analyses, we discovered a new bacterial phylum, *Ca.* Nitrosediminicolota. Metabolic potential analyses of *Ca.* Nitrosediminicolota genomes indicated that they contain the genetic machinery for nitrite oxidation, and also other versatile metabolisms such as nitrite reduction and urea utilization. *Ca.* Nitrosediminicolota is widespread in oligotrophic marine sediments. They are more abundant in the oxic zones of AMOR sediments by a factor of 2–4 compared to the canonical NOBs affiliated with *Nitrospiraceae* and *Nitrospinaceae*. Counting them as NOB resolves the abundance mismatch between AOA and NOB in marine oxic sediments and permits closing the nitrogen cycle in oxic marine sediments without invoking denitrification. Although being affiliated with the same genus, the two dominant *Ca.* Nitrosediminicola species in the AMOR sediments manifest distinct redox niche preferences: *Ca.* N. aerophilis is only present in the oxic zone whereas *Ca.* N. anaerotolerans exist both in the oxic and anoxic zones. Its capacity for thiosulfate reduction and amino acid fermentation may allow *Ca.* N. anaerotolerans to thrive under anaerobic conditions but cultivation efforts are needed to confirm these genome-based metabolic inferences. Considering their global occurrence and high abundance in sediments not only on the Arctic ridge but also beneath open ocean gyres and in hadal trenches, *Ca.* Nitrosediminicolota may play a critical role in sediment nitrogen cycling across the entire oligotrophic marine expanse.

### Etymology description

*Candidatus* Nitrosediminicolota (Ni.tro.se.di.mi.ni.co.lo’ta. N.L. masc. n. Nitrosediminicola, a bacterial genus; -ota, ending to denote a phylum; N.L. neut. pl. n. Nitrosediminicolota, the Nitrosediminicola phylum).

*Candidatus* Nitrosediminicola (Ni.tro.se.di.mi.ni’co.la. Gr. neut. n. nitron, mineral alkali; L. neut. n. sedimen, sediment; L. masc./fem. n. suff. -cola, inhabitant, dweller; N.L. masc. n. Nitrosediminicola, a nitrate forming sediment-dweller).

*Candidatus* Nitrosediminicola aerophilis (aero, oxygen; suff. -philis, lovers; aerophilis, an oxygen lover, highlighting the preference of this microbe to the oxic zone of marine sediments)

Phylogenetically affiliated with the genus *Ca.* Nitrosediminicola, phylum *Ca.* Nitrosediminicolota. This species currently contains three genomes from marine sediments (two from the Mariana Trench and one from the Arctic Mid-Ocean Ridge). The arctic genome consists of 73 scaffolds of 1,837,265 bp. The DNA G+C content is 60.6%. It is preferably present in the oxic sediment layers. It contains metabolic functions of aerobic nitrite oxidation and urea hydrolysis.

*Candidatus* Nitrosediminicola anaerotolerans (anaero, lack of oxygen; suff. -tolerans, being tolerant to something; anaerotolerans, being tolerant to anaerobic conditions, highlighting the tolerance of this microbe to the anoxic zone of sediment columns)

Phylogenetically affiliated with the genus *Ca.* Nitrosediminicola, phylum *Ca.* Nitrosediminicolota. This species contains two strains recovered from two Arctic sediment cores. Their genomes consist of 48–61 scaffolds, with total genome sizes of 2.2–2.5 Mbp. The DNA G+C content is 62.3%. The genomes are present in both the oxic and anoxic sediment layers. It contains metabolic functions of nitrite-nitrate conversion, urea hydrolysis, and thiosulfate reduction.

## Materials and Methods

### Sampling collection and characterization

This study uses samples and data generated and reported in ^28^ and ^14^, in which the procedures of sample collection, processing, and data generation were thoroughly described. Briefly, sediment cores were retrieved by gravity coring from the seabed of various sites on the ridge flanks of the Arctic Mid-Ocean Ridge beneath the Norwegian-Greenland Sea. The retrieved cores were split in halves for sampling and archiving. The thickness of the oxic zone of each core was determined by measuring the *in situ* oxygen concentrations using a needle-type fiber-optic oxygen microsensor (PreSens), except for GS13-CC2 in which the oxygen penetration depth was inferred as the depth marking the appearance of dissolved Mn in the porewater ^29^. The subsampling of microbiology samples (using sterile 10-mL cutoff syringes) and porewater extraction were performed immediately on the sampling half using Rhizons samplers after the split. Nitrate, nitrite, and ammonium concentrations in the porewater were measured colorimetrically by a Quaatro 114 continuous flow analyzer (SEAL Analytical Ltd).

### Exploring AOA and NOB abundances in marine oxic sediments

Similar to ^31^ where AOA’s distribution was explored, we investigated the distribution of NOB in oxic marine sediments based on the existing 16S rRNA gene amplicon sequencing data for 11 sediment cores with thick oxic zones. In addition to the cores considered in ^31^, we also included four additional AMOR cores (GS13-CC2, GS14-GC02, GS14-GC04, and GS15-GC01) reported in ^29, 30^ and two piston cores from the North Atlantic Gyre ^15^. The amplicon sequencing data of the total eight AMOR cores and the North Pond core were generated using the same procedure. Briefly, the total DNA in the sediment samples was extracted using the PowerLyze DNA extraction kits (MOBIO Laboratories, Inc.). Amplicon of the 16S rRNA gene was prepared using the two-round PCR amplification strategy with the “universal” primers of Uni519F/806r, as described in ^28^. The amplicon libraries were sequenced on an Ion Torrent Personal Genome Machine. As described in ^28^, sequencing reads were quality filtered and trimmed to 220 bp using the USEARCH v11.0.667 pipeline ^53^. The taxonomic classification of OTUs was performed using the lowest common ancestor algorithm implemented in the Python version of CREST4 (the latest version of CREST ^54^) against the SILVA 138.1 Release ^55^. The total cell numbers were taken as the sum of the archaeal and bacterial 16S rRNA genes as determined by qPCR. For the remaining two cores from the North Atlantic Gyre ^15^, we downloaded the amplicon sequencing data from the NCBI database and employed the same data analysis pipeline to run the reads trimming, OTU clustering and classification.

We initially considered the abundance of canonical NOB affiliated with the families *Nitrospiraceae* and *Nitrospinaceae*. For both AOA and canonical NOB, the absolute abundance of a functional group was calculated as the product of the total cell numbers (the sum of archaeal and bacterial 16S rRNA gene abundances) and its relative abundances in the total communities (as assessed by 16S rRNA gene amplicon sequencing), as previously employed for investigation of anammox bacteria ^28, 56^. We then considered also members of *Ca.* Nitrosediminicolota as some overlooked NOB in marine sediments and calculated the total NOB abundance by taking NOB abundance as the sum of *Nitrospiraceae*, *Nitrospinaceae*, and *Ca.* Nitrosediminicolota.

We also investigated the community structure of NOB based on the 16S rRNA gene amplicon sequencing data. Through the phylogenetic analysis of 16S rRNA gene sequences (see the description below), we confirmed that 10 OTUs were affiliated to the *Nitrospiraceae* family, 14 OTUs *Nitrospinaceae*, and 8 OTUs originally classified as members of the Schekmanbacteria phylum should correspond to the *Ca.* Nitrosediminicolota phylum. For each of these three groups, the relative abundance was taken as the sum of the relative abundances of the corresponding OTUs. To quantitatively evaluate the dominance of these three putative NOB lineages based on the observed depth profiles from arbitrarily selected sediment depths, we calculated the depth-averaged relative abundance for each of the three lineages using trapezoidal integration, as implemented in the R package *pracma* (https://github.com/cran/pracma).

### Genome binning and refinement

For metagenome-assembled genome recovery, we focused on the metagenome sequencing data of core GC08 and NP-1383E, which were generated and reported in ^28^. The procedures for DNA extraction, library preparation, metagenome sequencing, raw data quality control, assembly, and genome binning were described therein. Briefly, DNA was extracted from ∼7 g sediment of each selected depth. Metagenomic libraries were sequenced (2×150 bp paired-end) by an Illumina HiSeq 2500 sequencer. The quality of the raw sequencing data was first checked using FastQc v0.11.9 ^57^, with the adapters removed and reads trimmed using Trimmomatic v0.39 ^58^ based on the quality scores. The quality-controlled paired-end reads were *de novo* assembled into contigs using MEGAHIT v1.1.2 ^59^ with the *k*-mer length varying from 27 to 117. Contigs larger than 1000 bp were automatically grouped into genome bins using MaxBin2 v2.2.5 ^60^ and MetaBAT v2.15.3 ^61^ with the default settings, and the best representatives were selected using DAS_Tool v1.16 ^62^. The quality of the obtained bins was assessed using CheckM2 v1.0.2 ^63^. Only MAGs of ≥70% completeness with <5% redundancy were included for downstream analyses.

In this study, three NXR-containing MAGs (Bin_086, Bin_096, and Bin_108) were found to be affiliated with unknown bacterial phyla and were thus subject to further analyses. To ensure the binning correctness and also improve the MAG quality, quality-trimmed reads of the sample showing the highest genome coverage were mapped onto the contigs using BBmap ^64^, and the successfully aligned reads were re-assembled using SPAdes v3.12.0 ^65^ with the *k*-mers of 21, 33, 55, and 77. After the removal of contigs shorter than 1000 bp, the resulting scaffolds were visualized and manually re-binned using gbtools v2.6.0 ^66^ based on the GC content, taxonomic assignments, and differential coverages of contigs across multiple samples, with the input data generated using the following steps. Coverages of contigs in each sample were determined by mapping trimmed reads onto the contigs using BBMap v.37.61 ^64^. Taxonomic classification of contigs was assigned by BLASTn ^67^ according to the taxonomy of the single-copy marker genes in contigs. SSU rRNA sequences in contigs were identified using Barrnap ^68^ and classified using VSEARCH ^69^. The mapping, re-assembly, and re-binning process was repeated 5–7 times until the quality of the genomes could not be improved further. The refined MAGs were classified using GTDB-tk v2.3.0 ^70^ with the default setting. The MAG quality was checked again using CheckM2 v1.0.2 ^63^.

### Genome annotation

Genomes discussed in this study were annotated together with their close relative MAGs [i.e., three MAGs recovered from Mariana Trench sediments as reported in ^32^] and also representative MAGs in the phyla Nitrospinota and Nitrospinota_B in the GTDB 08-RS214 Release (https://gtdb.ecogenomic.org/). Genes in these genomes were predicted using Prodigal ^71^. Genome annotation was conducted using Prokka v1.13 ^72^, eggNOG ^73^, and BlastKoala ^74^ using the KEGG database. The functional assignments of genes of interest were also confirmed using BLASTp ^68^ against the NCBI RefSeq database. The metabolic pathways were reconstructed using KEGG Mapper ^75^.

### Linking MAGs with amplicon sequencing OTUs

To track the vertical distribution pattern of the two *Ca.* Nitrosediminicola species in the four AMOR cores, we searched the corresponding OTUs of the two genomes by comparing their 16S rRNA gene sequences (i.e., the query sequences) with the amplicon sequencing OTUs (the subject sequences) with BLASTp ^76^. Because Bin_096 reconstructed from AMOR sediments lacks a 16S rRNA gene sequence, we used that of B6D1T2 (another strain highly similar to Bin_096) to run the comparison. *Ca.* Nitrosediminicola Bin_086 has a full-length (1,565 bp) 16S rRNA gene sequence, which is a 100% match with OTU_4. *Ca.* Nitrosediminicola Bin_096 corresponds to OTU_235, given the 99.6% match of the 16S rRNA gene between them.

### Comparative genomic analysis

We performed a comparative analysis on the three representative genomes of the two *Ca.* Nitrosediminicola species using Anvi’o v7.1 ^77^ according to the pangenome analysis workflow. All genomes were first annotated using Prokka v.1.14 ^72^ and BLASTp using the Clusters of Orthologous Groups of Proteins (COG) ^78^ as the reference database. The comparative genomic analysis uses BLAST to quantify the similarity between each pair of genes, and the Markov Cluster algorithm (MCL) ^79^ (with an inflation parameter of 2) to resolve clusters of homologous genes. The shared and unique genes in the two genomes were identified via the functional enrichment analysis ^80^. Average amino acid identities between genomes were calculated using EzAAI v.1.2.2 ^81^ with the default setting.

### Phylogenetic analyses

To pinpoint the phylogenetic placement of the newly recovered MAGs and their relative genomes, we performed phylogenetic analyses for them together with high-quality genomes that were included in the GTDB Release 08-RS214. The 120 single-copy genes were identified, aligned, and concatenated using GTDB-tk v2.3.0 ^70^ with the “classify_wf” command. The maximum-likelihood phylogenetic tree was inferred based on this alignment using IQ-TREE v1.5.5 ^82^ with LG+F+R7 the best-fit model selected by ModelFinder ^83^, and 1000 ultrafast bootstrap iterations using UFBoot2 ^84^. To provide support to this phylogenomic analysis, we also performed the phylogenomic analysis based on the 14 syntenic ribosomal proteins (rpL2, 3, 4, 5, 6, 14, 16, 18, 22, and rpS3, 8, 10, 17, 19) that have been demonstrated to undergo limited lateral gene transfer ^85^. These selected proteins were identified in Anvi’o v7.1 ^77^ using Hidden Markov Model (HMM) profiles and aligned individually using MUSCLE ^86^. Alignment gaps were removed using trimAl ^87^ in “automated” mode. Individual alignments of ribosomal proteins were concatenated. The maximum likelihood phylogenetic tree was reconstructed using IQ-TREE v1.5.5 ^82^ with LG+R7 as the best-fit model.

A maximum-likelihood phylogenetic tree based on 16S rRNA genes was also constructed for the above-mentioned genomes to confirm the phylogenetic placement of the *Ca.* Nitrosediminicolota phylum. To expand this phylum on the tree beyond the available genomes, the putative *Ca.* Nitrosediminicolota OTUs from the amplicon sequencing and their close relatives identified via BLASTn ^76^ in the NCBI database were also included. Sequences were aligned using MAFFT-LINSi ^88^ and the maximum-likelihood phylogenetic tree was inferred as above, with 1,000 ultrafast bootstraps.

For the phylogenies of NxrA (nitrite oxidoreductase alpha subunit), the *Ca.* Nitrosediminicola sequences were used as the queries in BLASTp ^76^ searches in the NCBI database (>50% similarity and *E*-value of 10^-6^) to identify their close relatives. These sequences were aligned using MAFF-LINSi ^88^ with reference sequences from ref. ^89^, and complemented with known nitrite-oxidizing bacteria. For the phylogeny of UreC (urease alpha subunit), the sequences of *Ca.* Nitrosediminicola genomes were used as the queries in the BLASTp ^76^ search in the NCBI database (only hits of >50% similarity were retained), to identify their close relatives. These sequences were combined with sequences from ref. ^29^ and were aligned using MAFF-LINSi ^88^. The same procedure was also used to prepare sequences for the phylogenetic analyses of NirK (copper-containing nitrite reductase) and heme copper oxygen reductase. Phylogenetic trees for all proteins were generated as above.

### Global occurrence of *Ca.* Nitrosediminicolota

The global occurrence of *Ca.* Nitrosediminicolota in natural environments was assessed using IMNGS ^34^ against all public SRA datasets in the NCBI database with the 16S rRNA gene sequences of high-quality *Ca.* Nitrosediminicolota genomes as the query. Reads were counted as matching reads if they (1) were longer than 200 bp and (2) showed >95% nucleotide sequence identity to the query. Samples with less than 10 matching reads were discarded. Only natural environments with more than 0.1% relative abundances were retained for spatial mapping.

## Data availability

All sequencing data used in this study are available in the NCBI Short Reads Archive under the project number PRJNA529480. The three *Ca.* Nitrosediminicolota genomes recovered in this study are available under the accession number JAWJBM000000000 (Bin_086), JAWJBN000000000 (Bin_108), and JAWJBO000000000 (Bin_096).

## Acknowledgments

This work was funded by Simons Foundation grant 622065 and National Science Foundation grants OCE-2138890 and OCE-2142998 to A.R.B. R.Z. was supported by the MIT Molina Postdoctoral Fellowship. We are additionally grateful for the generosity of Dr. Bruce Heflinger in supporting the bablab, including this work. S.L.J was supported by the Trond Mohn Foundation and the University of Bergen through the Centre for Deep Sea research (grant TMS2020TMT13).

