## Supplementary Information for "A new abundant nitrite-oxidizing phylum in oligotrophic marine sediments"

**Table of Contents**

**Supplementary Text**

**Supplementary Figure S1–S7**

**Supplementary References**

**Supplementary Text**

**The capacity for urea hydrolysis in *Ca.* Nitrosediminocolota**

*Ca.* Nitrosediminocolota members can use urea as an alternative energy source. Similar to the NXR operon, the full urease operon (ureABCDFG) is present in the four *Ca.* Nitrosediminocolota genomes of higher completion (Fig. 3) and a partial urea operon in B13D1T1, while its absence in the remaining MAG (Bin_096) may be attributed to the low genomic completeness. It includes a nickel-dependent urease (UreABC) as well as accessory proteins (UreDFG) for the maturation of the holoenzyme. Additionally, all these urease-positive genomes possess a complete gene set for an ATP-dependent ABC-type urea transporter (UrtABCDE) encoded upstream of the urease structural genes. This type of transporter is characterized by its high affinity for urea, indicating an adaptation to low urea concentrations in the environment, such as marine sediments where tens of nanomolar concentrations of urea have been detected [1, 2]. In oxygenated environments where ammonium is maintained at low concentrations due to aerobic ammonia oxidation activity, urea can reach similar 10s to 100s of nanomolar levels as to ammonium [2]. In AMOR sediments, *Ca.* Nitrosediminocolota is the third phylum harboring microbes capable of urea utilization, after anammox bacteria affiliated to Planctomycetota [3, 4] and AOA affiliated to Thaumarchaeota [5].

*Ca.* Nitrosediminocolota bacteria may not oxidize ammonium directly because none of the five *Ca.* Nitrosediminocolota genomes has an ammonia monooxygenase. However, by providing a source of ammonium, the capacity of urea lysis of *Ca.* Nitrosediminocolota may allow reciprocal feeding with ammonia oxidizers [6]. In this substantial ecological advantage for NOBs [6], the ammonium released from urea degradation can serve as the substrate of the ammonia oxidizers, which in turn can provide nitrite to *Ca.* Nitrosediminocolota. This ecological advantage may be important to *Ca.* Nitrosediminocolota to grow in ammoniumlimited habitats such as the oxic zone of marine sediments. Urease is not universally present in NOB, and it is notably absent in the recently reported *Nitrospinaceae* and *Nitrospiraceae* NOB cultured from coastal sediments [7, 8]. The patchy distribution of urease among NOB suggests further niche differentiation of these organisms based on the capacity to use organic nitrogen compounds (e.g., urea) as sources of reduced nitrogen for assimilation.

Despite their close phylogenetic relationship, *Ca.* Nitrosediminocolota may acquire the urease via a route different from that of Nitrospinota. Phylogenetic analysis of UreC (urease alpha submit) indicates that *Ca.* Nitrosediminocolota genomes form a clade distinct from other bacterial phylum, and particularly other nitrogen cycling guilds (e.g., AOA, AOB, and NOB from the Nitrospirota and Nitrospinota phyla) (Fig. S5), suggesting that *Ca.* Nitrosediminocolota acquired urease differently from other nitrogen cycling groups. UreC sequences of *Ca.* Nitrosediminocola members show similarities to those of some Firmicutes (Fig. 3), indicating potential horizontal gene transfer (HGT) events between these two bacterial phyla given that these two phyla are not in close proximity on the tree of bacteria. It is likely that, like NXR, urease was horizontally disseminated between bacteria on multiple occasions.

**Similarity to other characterized NOBs**

Aerobic microbes display a variety of mechanisms, such as superoxide dismutase and catalase, to resist the oxidative stress caused by reactive oxygen species prevalent in oxic environments. While such canonical systems seem to be absent in some *Nitrospinaceae* genomes [e.g., [9, 10]], five of the six *Ca.* Nitrosediminocola genomes have superoxide dismutase (Fig. 3A), which may serve as a reactive oxygen species protection mechanism and help them to maintain anoxic niches in the oxic zone. Whether the presence of superoxide dismutase in *Ca.* Nitrosediminocola contribute to their ecological success in oxic AMOR sediments remains for further studies.

*Ca.* Nitrosediminocola members appear to lack the capacity for using formate, with formate dehydrogenase is only present in one of the six *Ca.* Nitrosediminocola MAGs. Formate dehydrogenase is present in most functionally characterized NOBs, and has been experimentally confirmed in *N. moscoviensis* [6, 11], *N. marina* [12], and *Nitrotoga fabula* [13], in which formate is used as an energy source and electron donor with nitrate as the terminal electron acceptor [6, 11, 12]. All six *Ca.* Nitrosediminocola genomes also lack the genes for flagellar synthesis and chemotaxis (Fig. 3), indicating that they may not be able to migrate within the sediment pore space, and thus are confined to the geochemical conditions of the depth at which they reside.

### Supplementary Figures

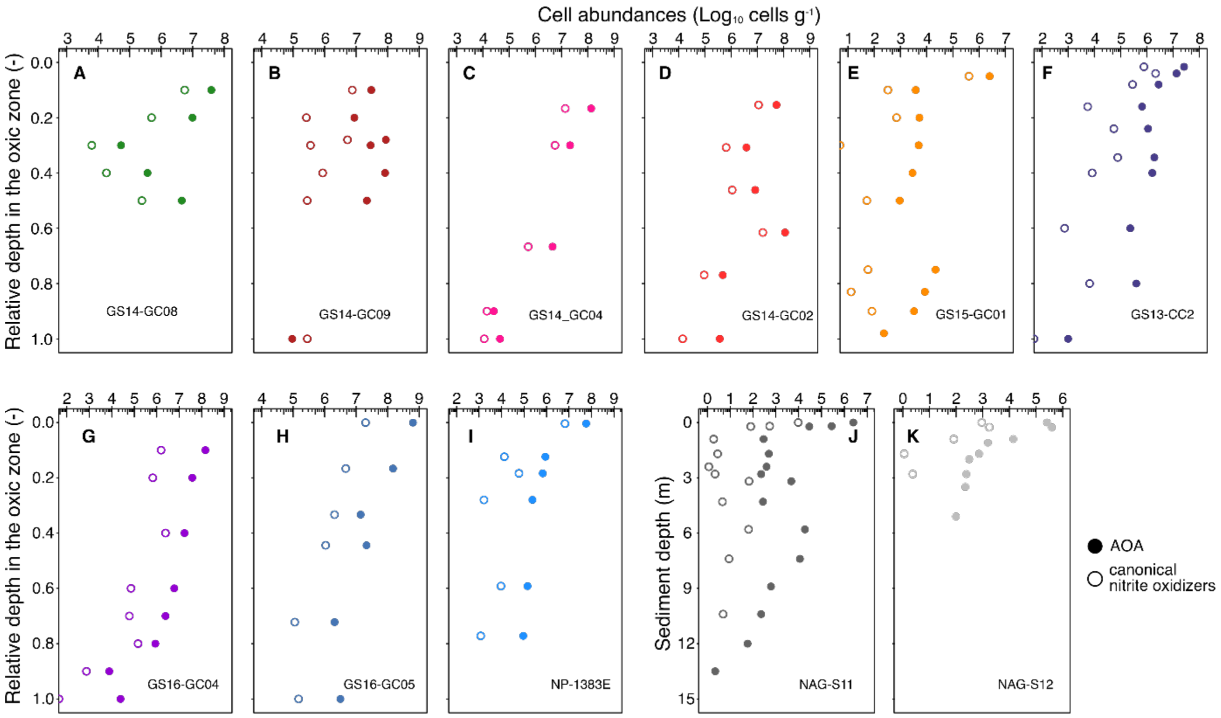


**Fig. S1. Absolute abundances of ammonia-oxidizing archaea (AOA) and canonical nitrite-oxidizing bacteria (the sum of *Nitrospiraceae* and *Nitrospinaceae*) in the oxic zones of eleven sediment cores with extensive oxic zones.** The abundances were calculated as the product of the total cell abundances and the relative abundances of the groups of interest in the total microbial communities as assessed by 16S rRNA gene amplicon sequencing. In cores **A**–**I**, relative depths within the oxic zone (0=sediment surface, 1=oxygen penetration depth) are shown to aid in comparison. Cores **J** and **K** are plotted against true depth below seafloor because the oxygen penetration depth was not resolved.

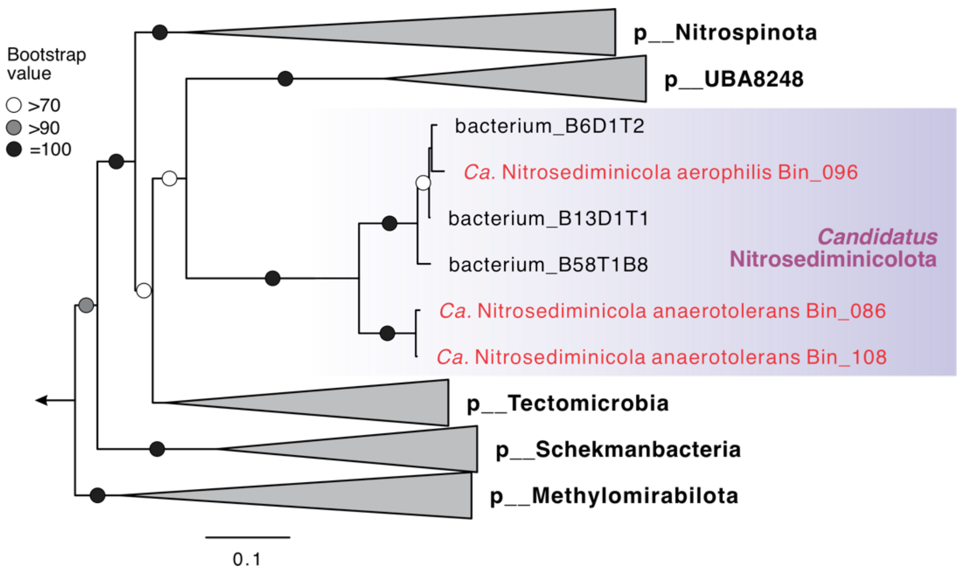


**Fig. S2. Maximum-likelihood phylogenetic tree of *Candidatus* Nitrosediminicolota and related bacterial phyla based on the concatenated 14 ribosomal proteins.** The tree is inferred using IQ-TREE with LG+R5 as the best-fit evolutionary model and 1,000 ultrafast bootstrap iterations. The nomenclature of the bacterial phyla follows GTDB, except that *Candidatus* Nitrosediminicolota is proposed in this study.

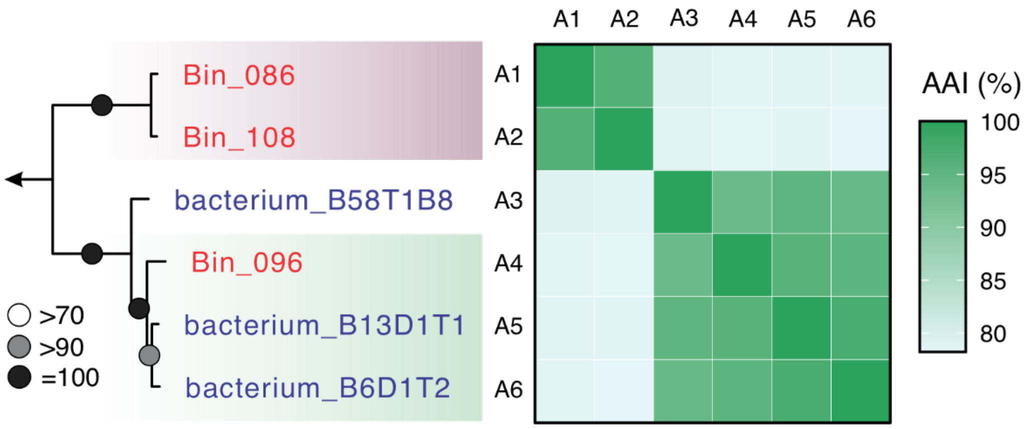


**Fig. S3. Average amino acid identity (AAI) between genomes in the newly proposed phylum *Candidatus* Nitrosediminicolota.** The three MAGs recovered in this study are highlighted in red, whereas those pre-existing from previous studies are in blue. The phylogenetic tree shown on the left side was a subset of Fig. 2A. The lineage with the red background is *Ca.* N. anaerotolerans, while the one with light green background represents *Ca.* N. aerophilis.

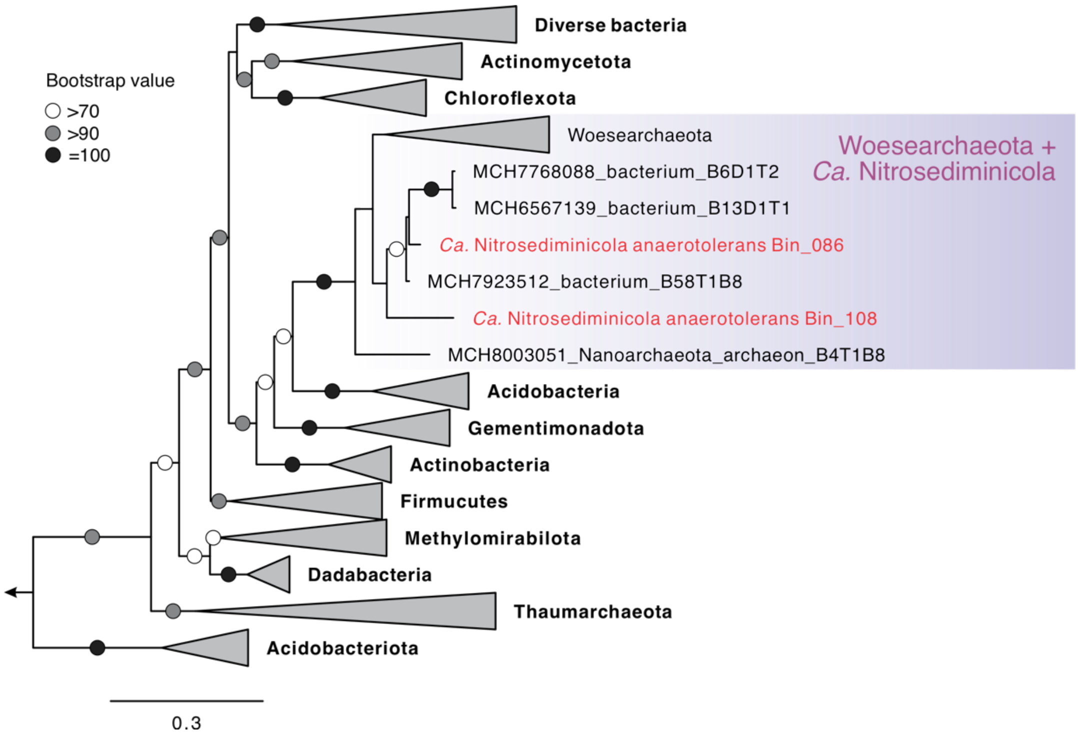


**Fig. S4. Maximum-likelihood phylogenetic tree of copper-containing nitrite reductase (NirK).** All sequences included here belong to "Clade II" of NirK. Only sequences of *Ca.* Nitrosediminicola described in this study are shown, while sequences of other various bacterial phyla are collapsed. The tight association of NirK sequences between *Ca.* Nitrosediminicola and the archaeal lineage *Ca.* Woesearchaeota is highlighted by a colored box.

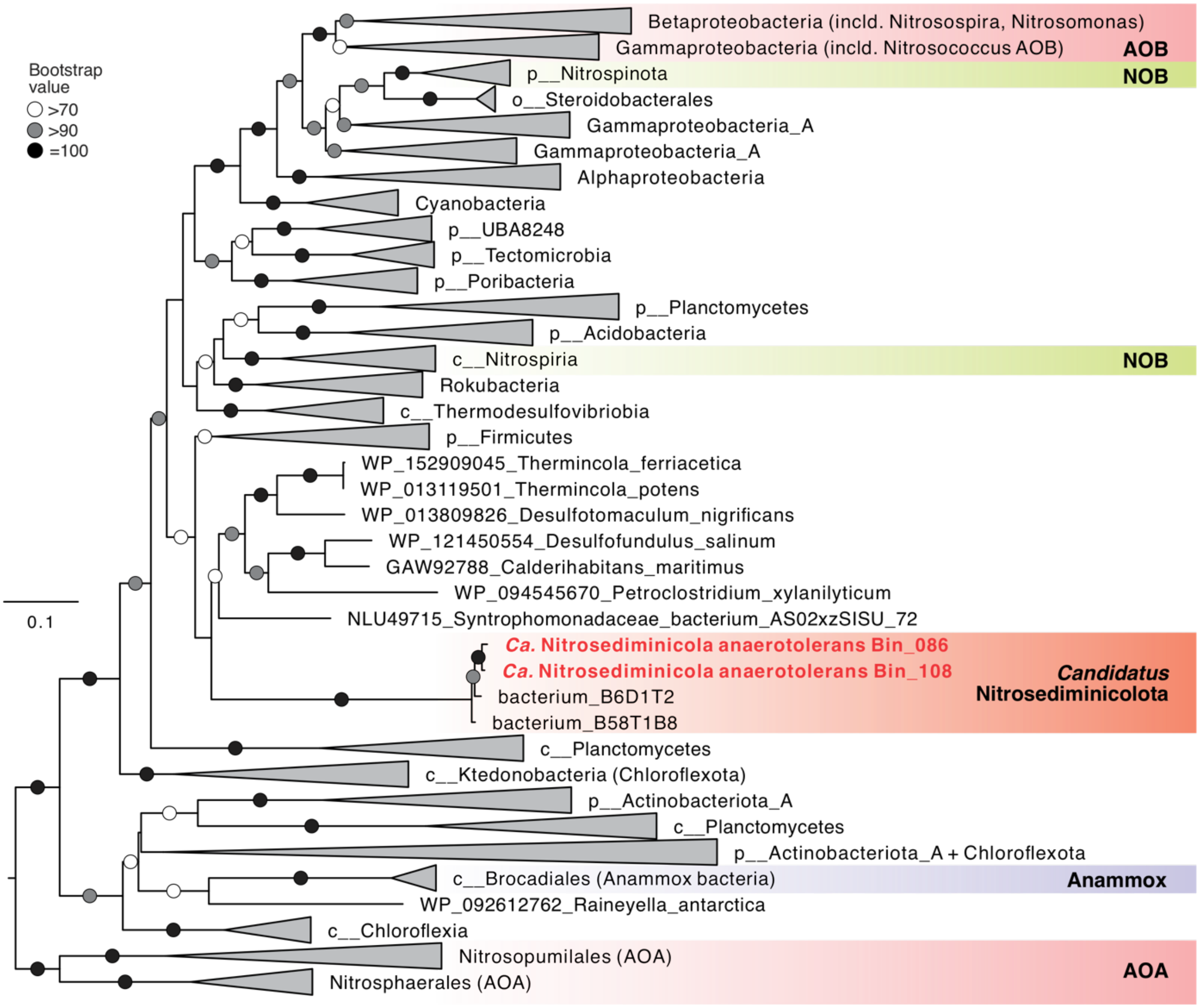


**Figure S5. Maximum-likelihood phylogenetic tree of UreC (urease alpha subunit).** The phylogeny was reconstructed using IQ-tree v1.6.10 under the LG+C20+F+G substitution model with 1,000 ultrafast bootstraps. The tree was rooted to NarG sequences of NC10 bacteria. Genomes recovered in this study are shown in red. Bacteria known for having the capacity of nitrite oxidation (i.e., nitrite-oxidizing bacteria of the genera of *Nitrospira*, *Nitrospina*, *Nitrotoga*, *Nitrobacter*, and *Nitrococcus*, and anammox bacteria of the Brocadiales order) are highlighted in orange. Bacteria with an observed nitrate-reducing phenotype are shown in purple. Bootstrap values >70 are shown with symbols listed in the legend. The scale bar shows estimated sequence substitutions per residue.

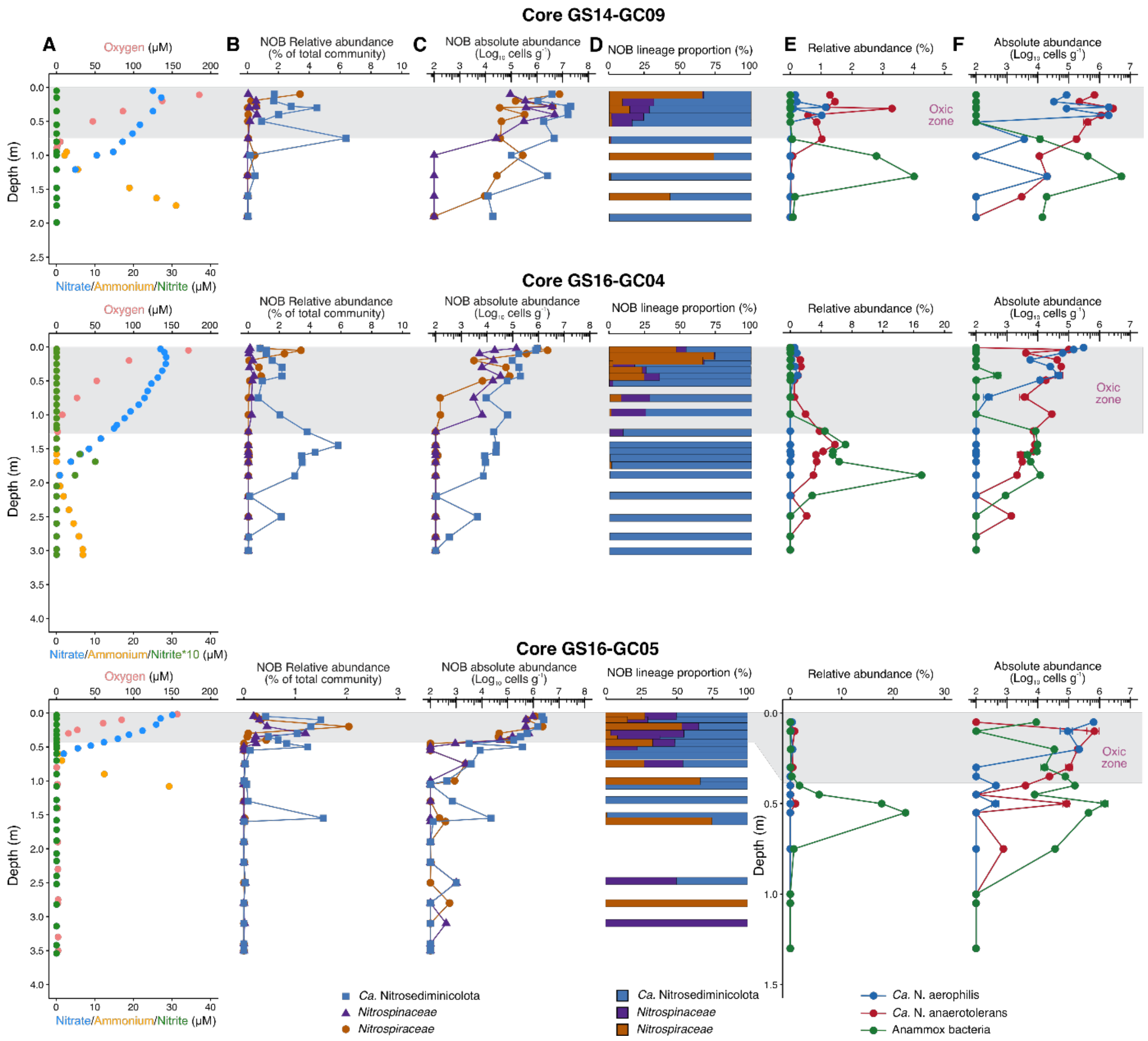


**Fig. S6. Geochemical context, relative abundances, communication compositions of NOB lineages in AMOR cores GS14-GC09, GS16-GC04, and GS16-GC05. (A)** Geochemical context delineated by the measured profiles of oxygen, nitrate, nitrite, and ammonium, previously reported in Zhao et al. (2020). The oxic zone in each core is marked with a grey box. **(B)** The relative abundances of *Ca.* Nitrosediminicolota and the canonical marine NOB families *Nitrospiraceae* and *Nitrospinaceae*, as assessed by amplicon sequencing. **(C)** The absolute abundances of the three NOB lineages calculated as the product of the relative abundances of the three lineages and the total cell numbers. **(D)** The community composition of NOB community in each of the investigated depth. **(E, F)** The relative **(E)** and absolute **(F)** abundances of two *Ca.* Nitrosediminicola species (*Ca.* N. aerophilis and *Ca.* N. anaerotolerans) and anammox bacteria throughout the core. Note the last two panels in core GS16-GC05 are shown in a small vertical scale to highlight the variations in the upper sediment layers.

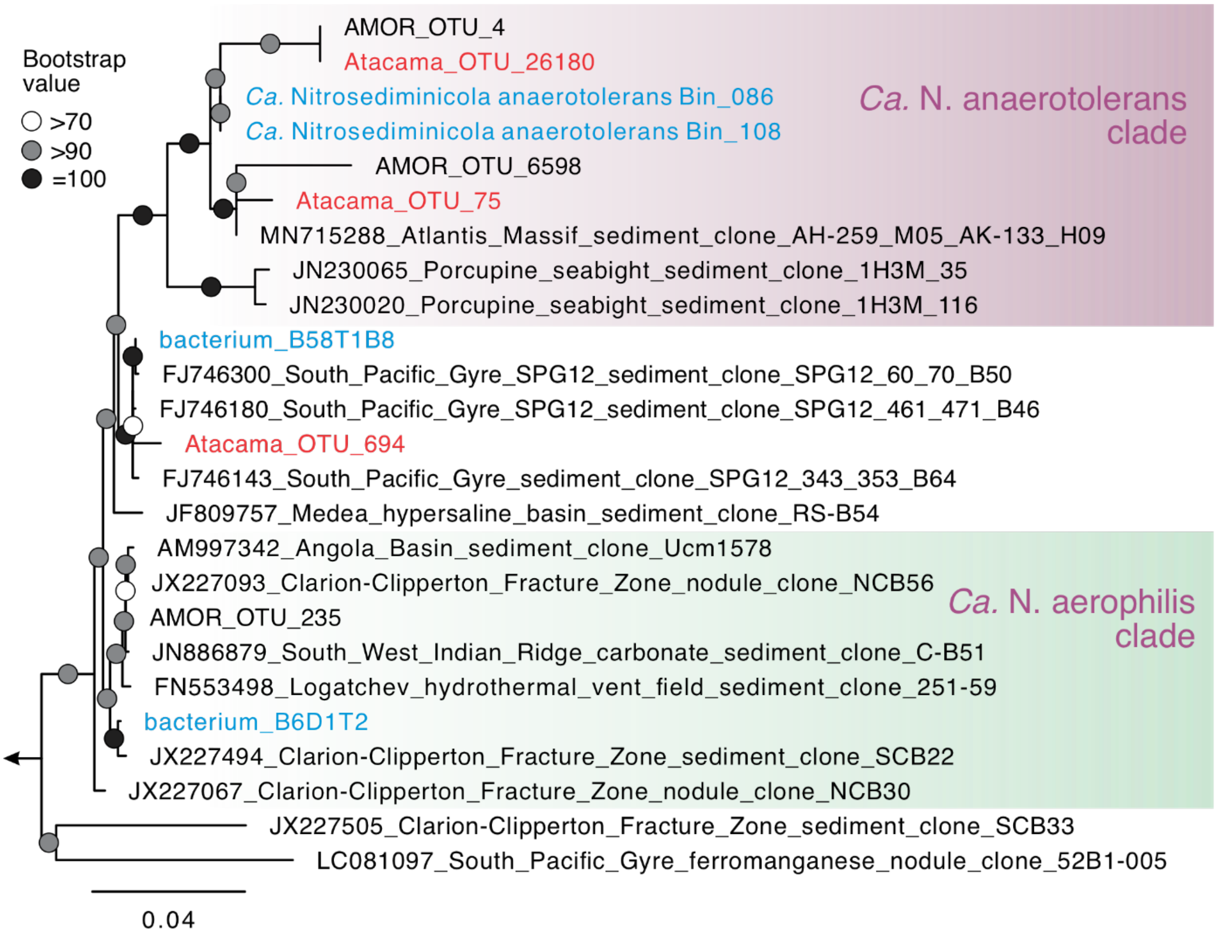


**Fig. S7. Phylogenetic affiliation of the *Ca.* Nitrosediminicolota OTUs in Atacama cores.** The three OTUs from the Atacama sediment cores are highlight in red, and the clades emcompassing *Ca.* N. anaerotoletans and *Ca.* N. aerophilis are distinguished by red and green backgrounds, respectively.
